# Do altered neural responses during self-reflection converge across psychiatric disorders? A systematic review and meta-analysis of neuroimaging studies

**DOI:** 10.64898/2026.03.13.711269

**Authors:** Shanshan Zhu, Wen-Jing Yan, Hu Chuan-Peng

## Abstract

Altered neural responses during self-reflection have been reported across psychiatric disorders, but it is unknown whether there exist common brain regions across studies. To fill the gap, we conducted a systematic review and coordinate-based activation likelihood estimation (ALE) meta-analysis of task-based neuroimaging studies. Through systematic literature search and article screening, we identified 36 studies across 8 diagnostic categories, of which 25 studies were eligible for the ALE meta-analysis. Qualitatively, these studies revealed altered neural response among psychiatric disorders in brain regions include cortical midline and lateral association regions. ALE analyses were conducted separately for self-versus other and self-versus control contrasts, the former identified convergence in the right anterior cingulate cortex (ACC), and the latter found converged activation in the left middle frontal gyrus (MFG). These findings revealed a broad interest in neural response to self-evaluation but unevenly distributed studies across psychiatric disorders, and the available results only provided weak evidence of spatial convergence for altered neural response across psychiatric disorders.

## 1 Introduction

Self-referential processing is an umbrella term referring to a variety of mental processes that support the self-concept (Hards et al., 2024). These processes are essential for individual identity, mental health, and social functions (Leary & Tangney, 2012; Northoff, 2013; Northoff et al., 2026) and are a core construct in cognitive science, psychology, and psychiatry (Gallagher, 2000; Nelson et al., 2014; Northoff et al., 2006; Philippi & Koenigs, 2014). Among these self-referential processes, self-reflection, an explicit evaluation of one’s own psychological characteristics, is of particular interest in mental health because it involves individuals’ knowledge and beliefs about themselves (Hards et al., 2024). Thus, self-reflection is closely linked to people’s psychological or mental self (James, 1890). In the literature, self-reflection is used interchangeably with self-referential (van der Meer et al., 2010), partially because the neural basis of self-reflection is typically studied with a behavioral task known as the “self-referential encoding task” (e.g., Kelley et al., 2002; Zhang et al., 2006), in which participants first reflect on whether trait words describe themselves or others and are then asked to perform a memory task (Rogers et al., 1977; Sun et al., 2023).

Disturbances in self-concept as revealed by self-reflection are widely reported across major psychiatric disorders and are increasingly considered central to psychopathology. For example, disturbances of the sense of self are widely reported in schizophrenia, including fragmentation of self-experience, and loss of ego boundaries (Raballo et al., 2025). Similarly, individuals with major depressive disorder often exhibit persistent negative self-evaluations (Fusar-Poli et al., 2023). More often than not, a clearer self-concept is an important goal of individuals who seek psychotherapy (Ladmanová et al., 2025). These findings suggest that aberrant self-concept as reflected in self-reflection is not confined to a single disorder but represents a pervasive feature across psychiatric conditions.

Indeed, behavioral evidence supports its transdiagnostic relevance. Most content of repetitive negative thinking is related to reflecting one’s own experience and shaping one’s self-concept (Moulds & McEvoy, 2025). The discrepancies between actual and ideal self-representations, which are often the result of self-reflection, have been proposed as transdiagnostic features (Mason et al., 2019). However, direct evidence for the transdiagnostic significance of the neural mechanisms underlying such aberrations remains scarce. While meta-analyses of resting-state fMRI studies reported alterations in default mode network (DMN) connectivity across psychiatric disorders (Doucet et al., 2020), interpreting these findings as altered self-reflection is speculative because the DMN involved in multiple psychological processes (Menon, 2023). Task-based fMRI studies, which contrasted neural responses during self-versus other conditions, provided more direct evidence for self-reflection (Potvin et al., 2019). Consistent with this notion, recent systematic reviews reported shared altered neural responses in midline and lateral cortical regions across schizophrenia, depression, and anxiety (Sabbah & Northoff, 2024, 2025). However, these reviews did not distinguish different self-referential processes, which had been reported to be supported by different neural systems (Hu et al., 2016; Northoff et al., 2026; Qin et al., 2020). Thus, it remains unknown whether the changed neural response during self-reflection is shared across psychiatric disorders.

To address this question, we conducted a systematic review and meta-analysis to examine neural responses during self-reflection across psychiatric disorders. Drawing on theoretical discussions of transdiagnostic research (Barch, 2020; Dalgleish & Hitchcock, 2023), we considered the diagnostic breadth and direction of reported alterations when interpreting their potential transdiagnostic relevance. First, a putative transdiagnostic neural feature should be observed across multiple diagnostic categories. Second, the directionality of neural activations provides additional information about the nature of potentially shared regional involvement. For example, McTeague and colleagues (2020) reported that activations in the dorsomedial thalamus exhibited opposite patterns across disorders, with lower activation in psychotic disorders (specifically schizophrenia) but higher activation in nonpsychotic disorders such as anxiety and depression.

We systematically examined whether altered neural responses during self-reflection show spatial convergence across psychiatric disorders. Building on the considerations described above, we combined a PRISMA-guided systematic review with a coordinate-based activation likelihood estimation (ALE) meta-analysis. The systematic review synthesized existing neuroimaging evidence on self-reflection across psychiatric diagnoses, distinguishing the “self versus other” contrast from the “self versus control” contrast and characterizing the diagnostic coverage and direction of reported patient–control differences. We then conducted quantitative ALE analyses to test for spatial convergence of patient–control differences within each task-contrast category. By integrating systematic review methodology with complementary ALE approaches, this study aimed to evaluate the available evidence for potentially shared regional alterations during self-reflection under our framework and provide insight for future studies.

## 2 Methods

### 2.1 Literature search and article screening

We conducted a systematic review and meta-analysis of neuroimaging studies employing the self-referential encoding task in psychiatric disorders, following the Preferred Reporting Items for Systematic Reviews and Meta-Analyses (PRISMA) guidelines (Page et al., 2021). The activation likelihood estimation (ALE) meta-analysis procedure followed established recommendations for neuroimaging meta-analyses (Müller et al., 2018; Liu et al., 2021) and was preregistered on the Open Science Framework (OSF; registration identifier: https://osf.io/5bg9v/). Any deviations from the preregistered protocol were documented according to Willroth & Atherton (2024) and are reported in Supplementary Table S2.

Reference lists of two relevant systematic reviews were also screened (Sabbah & Northoff, 2024, 2025). The database searched and the keywords used are listed in Supplementary Table S1. EndNote 21 was used to manage the items and remove duplicates. After duplicates were removed, two independent reviewers identified the most pertinent articles based on the inclusion criteria:

1. Empirical research;
2. Papers written in English and formally published in academic journals;
3. The average age of participants ranged from 18 to 60 years old;
4. Studies were required to include participants with psychiatric disorders diagnosed. through standardised diagnostic criteria (including DSM-IV-TR, DSM-IV, ICD-10, MINI-Plus, and Autism Diagnostic Observation Schedule, module 4, Lord et al., 2018);
5. Functional magnetic resonance imaging (fMRI) studies or positron emission tomography (PET);
6. Experimental tasks should include at least one eligible contrast between self versus other (e.g., self>mother) and self versus control conditions (e.g., self>syllable judgement);
7. Reported peak coordinates in standard brain space (Talairach or MNI space).

Any conflicting decisions in the screening process were resolved by the senior author. Two independent coders performed the data extraction and coding procedures. Inter-rater reliability was assessed using the AC_1_ coefficient (Gwet, 2008). The AC_1_ coefficient for the data coding was 0.888, exceeding the 0.8 threshold and demonstrating a high degree of inter-rater consistency for the literature coding in this dataset. Detailed procedures of the reliability analysis are provided in the Supplementary Materials S2. The literature search and screening process is summarised in a PRISMA flow diagram (Figure 1).

**Figure 1.**
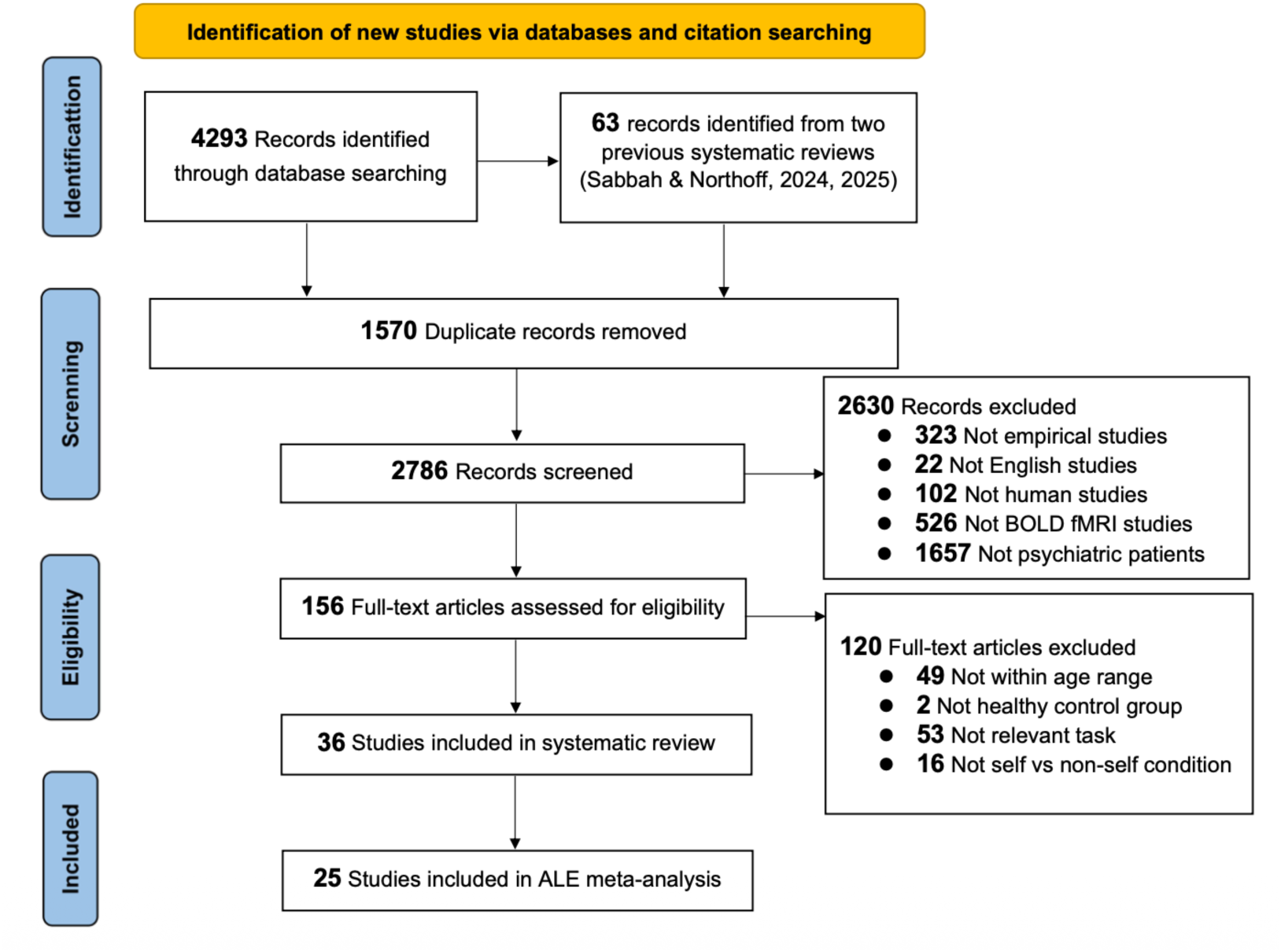
Preferred reporting items for Meta-analyses (PRISMA) flowchart detailing screening process.

### 2.2 Data extraction

To enable a systematic and quantitative analysis, we extracted key information from each included study. The extracted variables included study characteristics, participant demographics, imaging modality, and stereotaxic activation coordinates. For psychiatric patients, we additionally recorded the diagnostic category, diagnostic instrument, and medication status. Our comprehensive dataset is publicly available via the Scientific Data Bank (https://doi.org/10.57760/sciencedb.j00001.00469). In line with Müller et al. (2018), we distinguished between a “study” (an individual publication) and an “experiment” (a specific contrast yielding a set of peak coordinates, e.g., self-versus other-conditions).

Study characteristic included publication year, country and recruitment setting. Demographic and clinical variables included psychiatric diagnosis, diagnostic criteria and instruments, clinical status and scores, illness duration, comorbidities, medication status, and treatment information. Sample size, age, and sex were extracted separately for psychiatric and healthy control groups.

For each eligible task, we extracted the stimulus materials, presentation modality, judgment instructions, judgment targets, comparison conditions, and exact contrast reported. Self-reflection was operationalized as explicitly judging whether traits or statements applied to oneself. Task contrasts were classified as self versus other (e.g., “Does this trait word describe you or your mother?”) or self versus control (e.g., “Judging whether an attribute describes oneself versus counting the number of letters in a word such as ‘red’”). For experiments included in the ALE analyses, stimulus materials were linguistic or verbal, including words, sentences, or statements presented visually or auditorily. Task information and group sample sizes are provided in Table 1.

**Table 1.**
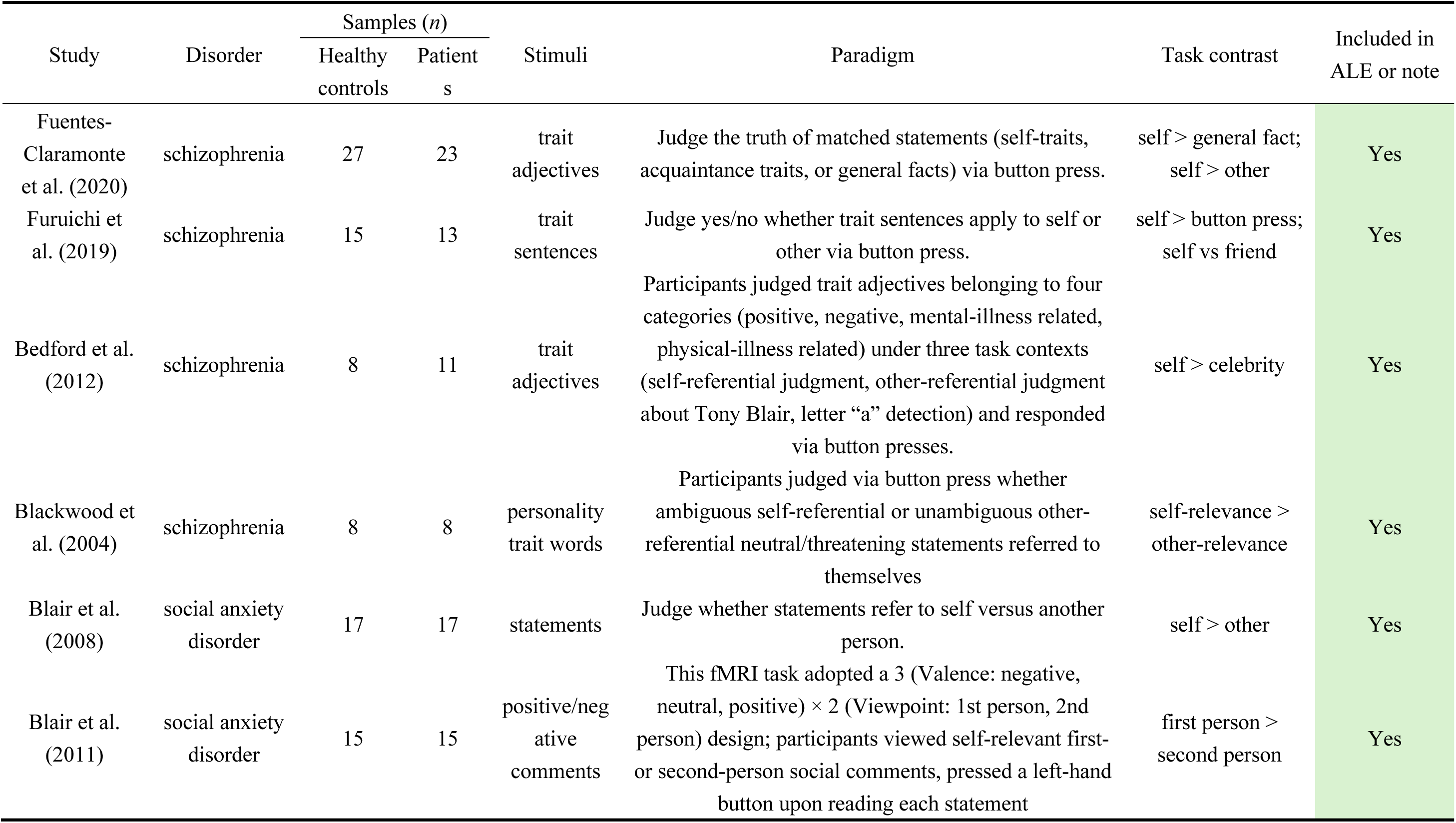
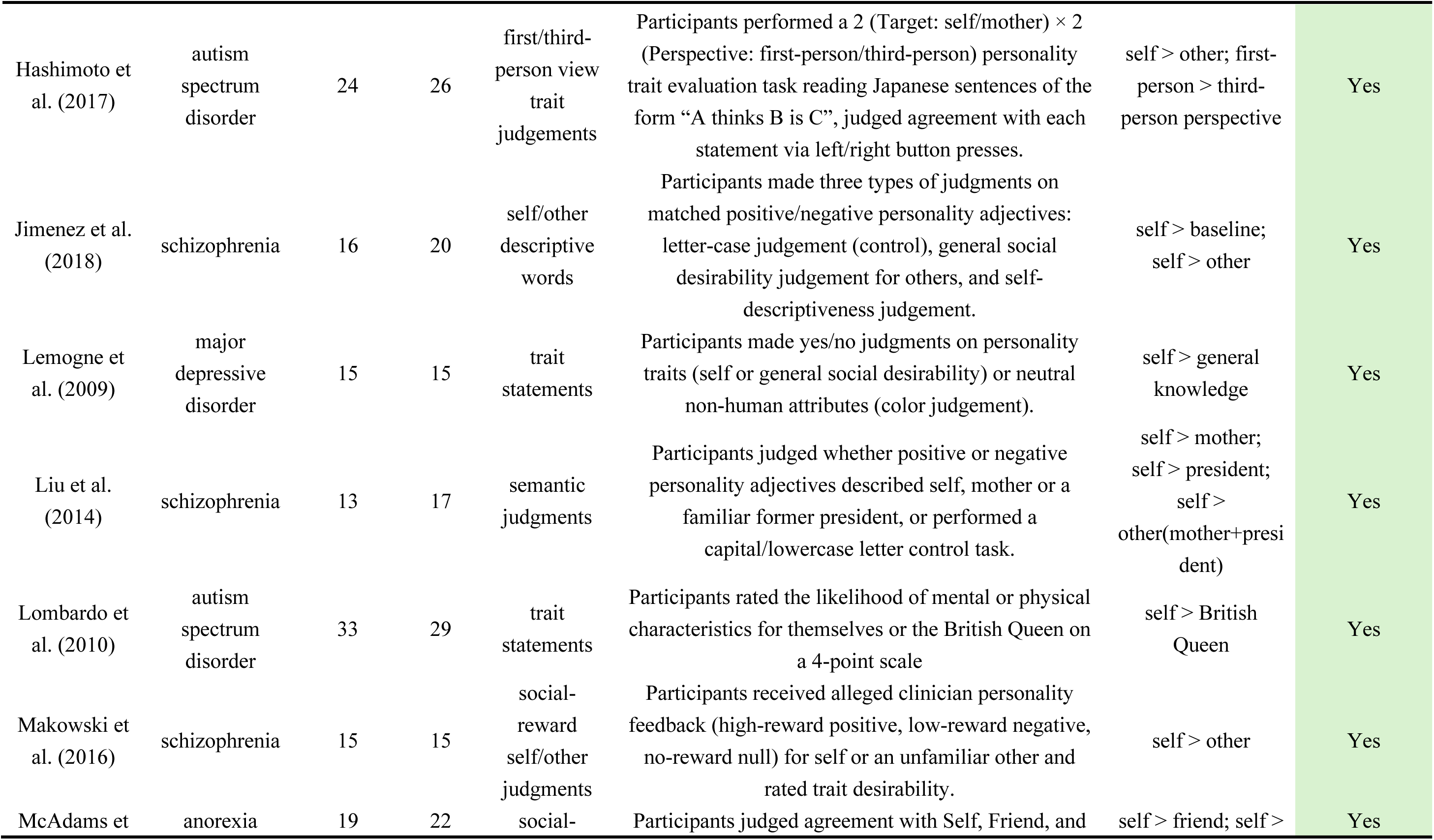
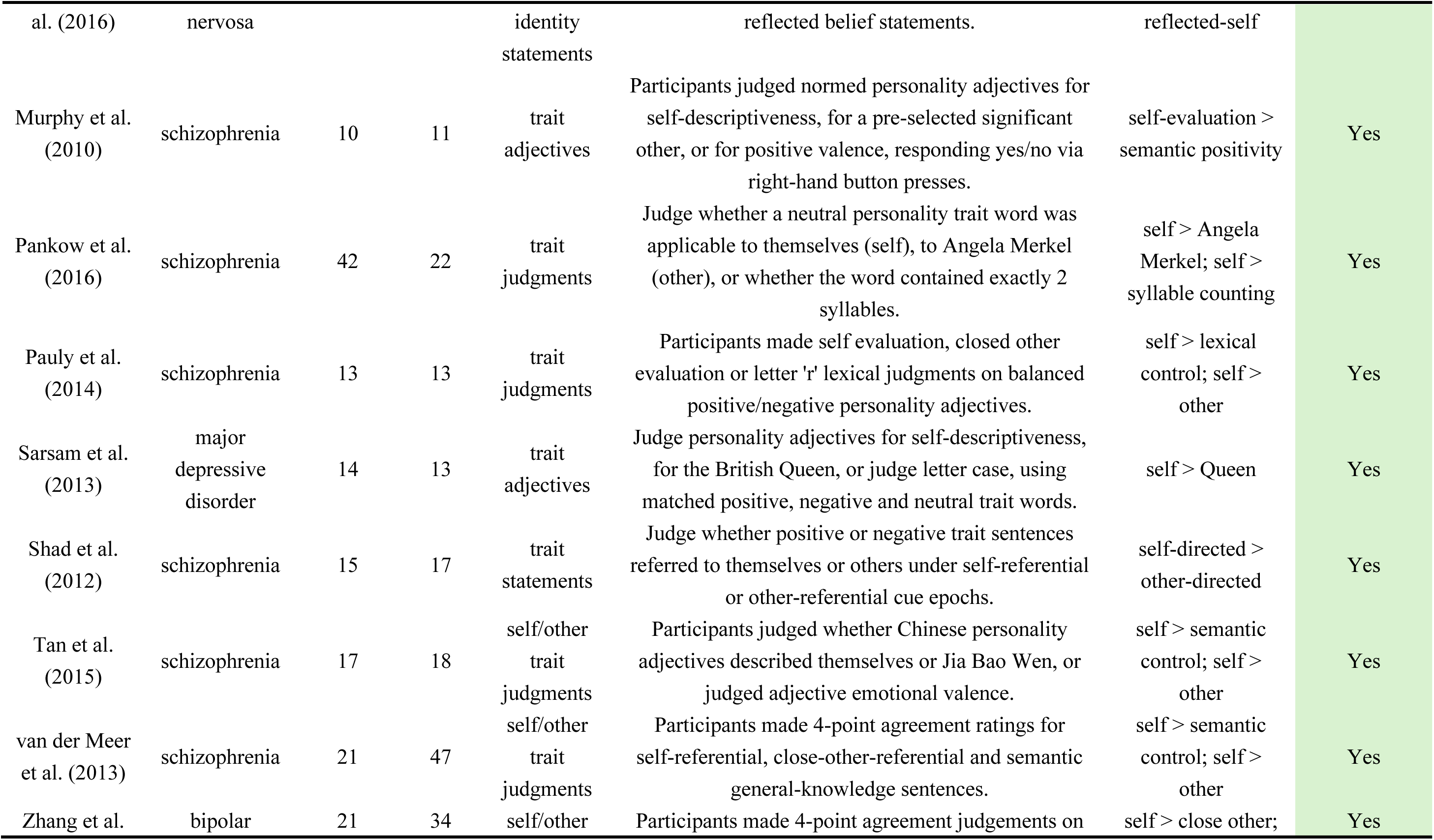
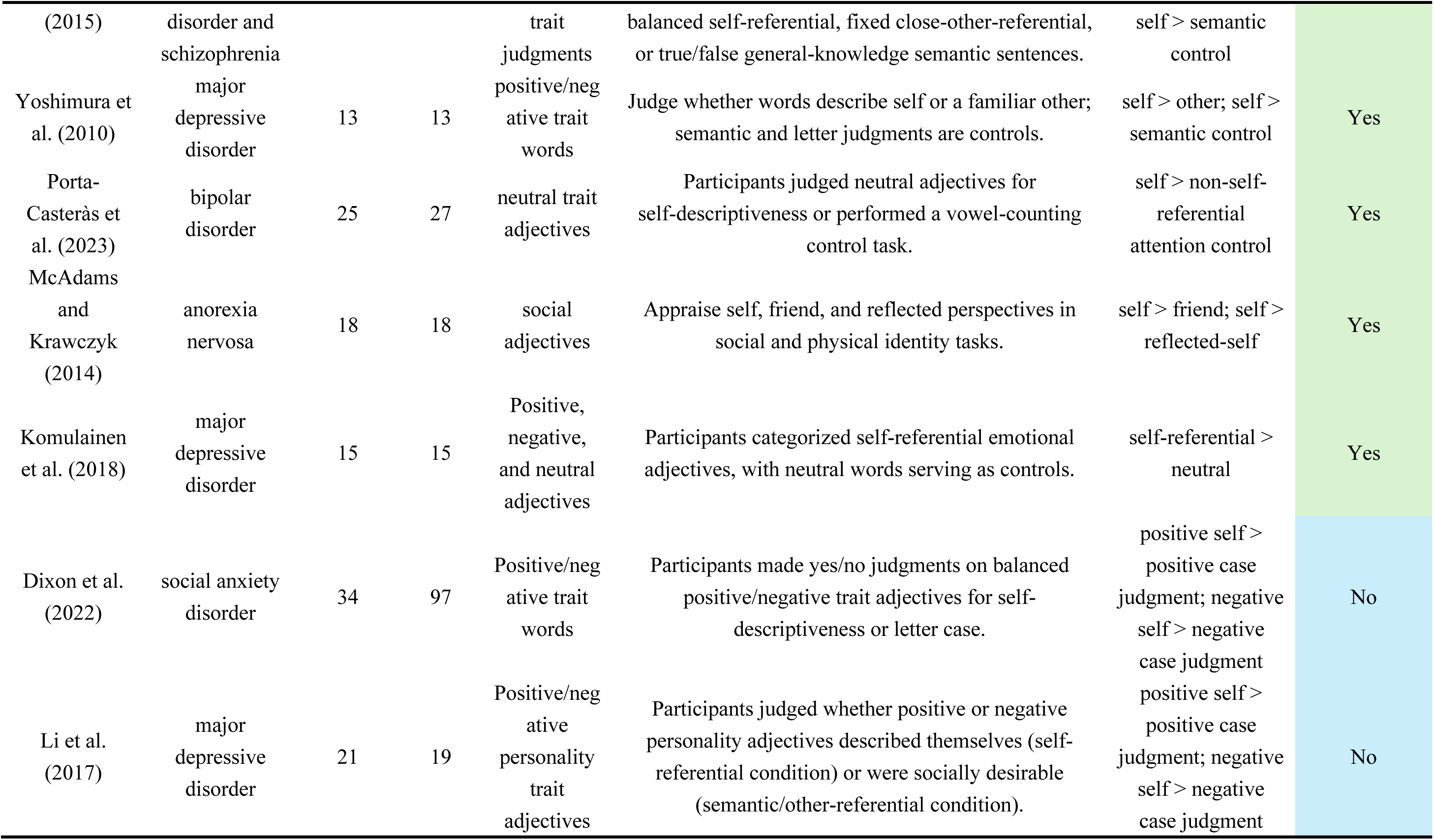
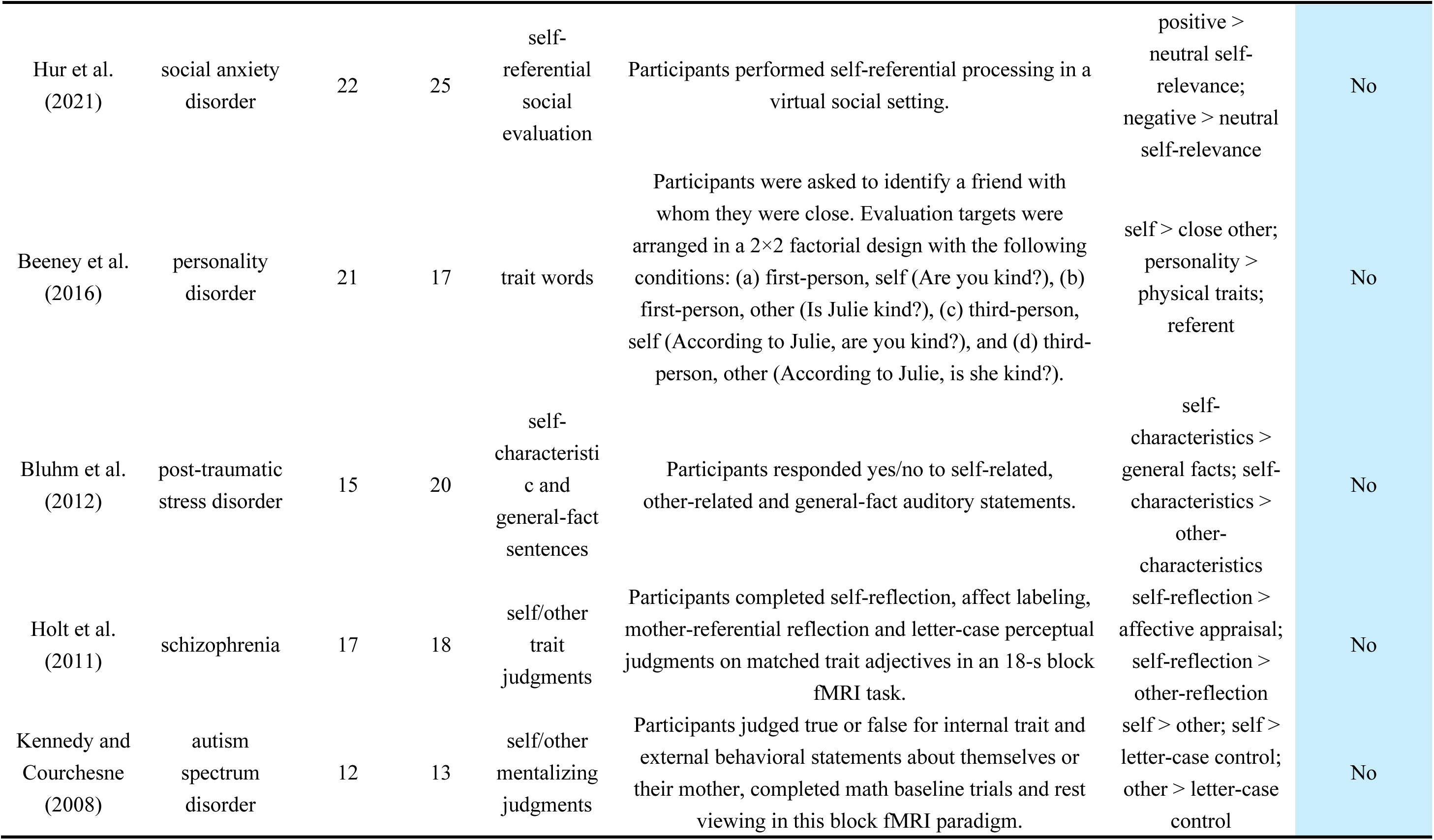
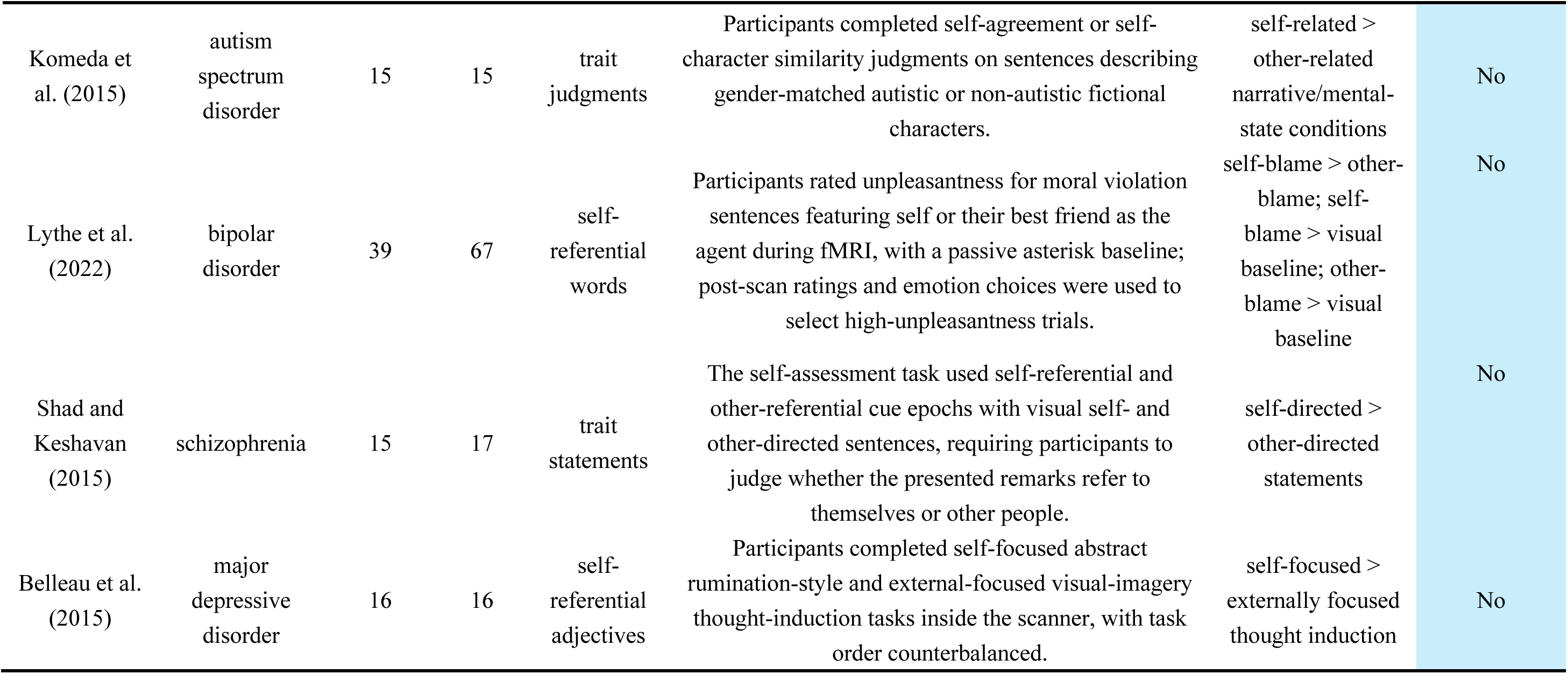
Study characteristics and task paradigms in the systematic review and meta-analyses (*n* = 36).

| Study | Disorder | Samples (*n*) |  | Stimuli | Paradigm | Task contrast | Included in ALE or note |
|---|---|---|---|---|---|---|---|
|  |  | Healthy controls | Patients |  |  |  |  |
| Fuentes-Claramonte et al. (2020) | schizophrenia | 27 | 23 | trait adjectives | Judge the truth of matched statements (self-traits, acquaintance traits, or general facts) via button press. | self > general fact; self > other | Yes |
| Furuichi et al. (2019) | schizophrenia | 15 | 13 | trait sentences | Judge yes/no whether trait sentences apply to self or other via button press. | self > button press; self vs friend | Yes |
| Bedford et al. (2012) | schizophrenia | 8 | 11 | trait adjectives | Participants judged trait adjectives belonging to four categories (positive, negative, mental-illness related, physical-illness related) under three task contexts (self-referential judgment, other-referential judgment about Tony Blair, letter “a” detection) and responded via button presses. | self > celebrity | Yes |
| Blackwood et al. (2004) | schizophrenia | 8 | 8 | personality trait words | Participants judged via button press whether ambiguous self-referential or unambiguous other-referential neutral/threatening statements referred to themselves | self-relevance > other-relevance | Yes |
| Blair et al. (2008) | social anxiety disorder | 17 | 17 | statements | Judge whether statements refer to self versus another person. | self > other | Yes |
| Blair et al. (2011) | social anxiety disorder | 15 | 15 | positive/negative comments | This fMRI task adopted a 3 (Valence: negative, neutral, positive) × 2 (Viewpoint: 1st person, 2nd person) design; participants viewed self-relevant first- or second-person social comments, pressed a left-hand button upon reading each statement | first person > second person | Yes |
| Hashimoto et al. (2017) | autism spectrum disorder | 24 | 26 | first/third-person view trait judgements | Participants performed a 2 (Target: self/mother) × 2 (Perspective: first-person/third-person) personality trait evaluation task reading Japanese sentences of the form “A thinks B is C”, judged agreement with each statement via left/right button presses. | self > other; first-person > third-person perspective | Yes |
| Jimenez et al. (2018) | schizophrenia | 16 | 20 | self/other descriptive words | Participants made three types of judgments on matched positive/negative personality adjectives: letter-case judgement (control), general social desirability judgement for others, and self-descriptiveness judgement. | self > baseline; self > other | Yes |
| Lemogne et al. (2009) | major depressive disorder | 15 | 15 | trait statements | Participants made yes/no judgments on personality traits (self or general social desirability) or neutral non-human attributes (color judgement). | self > general knowledge | Yes |
| Liu et al. (2014) | schizophrenia | 13 | 17 | semantic judgments | Participants judged whether positive or negative personality adjectives described self, mother or a familiar former president, or performed a capital/lowercase letter control task. | self > mother; self > president; self > other(mother+president) | Yes |
| Lombardo et al. (2010) | autism spectrum disorder | 33 | 29 | trait statements | Participants rated the likelihood of mental or physical characteristics for themselves or the British Queen on a 4-point scale | self > British Queen | Yes |
| Makowski et al. (2016) | schizophrenia | 15 | 15 | social-reward self/other judgments | Participants received alleged clinician personality feedback (high-reward positive, low-reward negative, no-reward null) for self or an unfamiliar other and rated trait desirability. | self > other | Yes |
| McAdams et al. (2016) | anorexia nervosa | 19 | 22 | social-identity statements | Participants judged agreement with Self, Friend, and reflected belief statements. | self > friend; self > reflected-self | Yes |
| Murphy et al. (2010) | schizophrenia | 10 | 11 | trait adjectives | Participants judged normed personality adjectives for self-descriptiveness, for a pre-selected significant other, or for positive valence, responding yes/no via right-hand button presses. | self-evaluation > semantic positivity | Yes |
| Pankow et al. (2016) | schizophrenia | 42 | 22 | trait judgments | Judge whether a neutral personality trait word was applicable to themselves (self), to Angela Merkel (other), or whether the word contained exactly 2 syllables. | self > Angela Merkel; self > syllable counting | Yes |
| Pauly et al. (2014) | schizophrenia | 13 | 13 | trait judgments | Participants made self evaluation, closed other evaluation or letter 'r' lexical judgments on balanced positive/negative personality adjectives. | self > lexical control; self > other | Yes |
| Sarsam et al. (2013) | major depressive disorder | 14 | 13 | trait adjectives | Judge personality adjectives for self-descriptiveness, for the British Queen, or judge letter case, using matched positive, negative and neutral trait words. | self > Queen | Yes |
| Shad et al. (2012) | schizophrenia | 15 | 17 | trait statements | Judge whether positive or negative trait sentences referred to themselves or others under self-referential or other-referential cue epochs. | self-directed > other-directed | Yes |
| Tan et al. (2015) | schizophrenia | 17 | 18 | self/other trait judgments | Participants judged whether Chinese personality adjectives described themselves or Jia Bao Wen, or judged adjective emotional valence. | self > semantic control; self > other | Yes |
| van der Meer et al. (2013) | schizophrenia | 21 | 47 | self/other trait judgments | Participants made 4-point agreement ratings for self-referential, close-other-referential and semantic general-knowledge sentences. | self > semantic control; self > other | Yes |
| Zhang et al. (2015) | bipolar disorder and schizophrenia | 21 | 34 | self/other trait judgments | Participants made 4-point agreement judgements on balanced self-referential, fixed close-other-referential, or true/false general-knowledge semantic sentences. | self > close other; self > semantic control | Yes |
| Yoshimura et al. (2010) | major depressive disorder | 13 | 13 | positive/negative trait words | Judge whether words describe self or a familiar other; semantic and letter judgments are controls. | self > other; self > semantic control | Yes |
| Porta-Casteràs et al. (2023) | bipolar disorder | 25 | 27 | neutral trait adjectives | Participants judged neutral adjectives for self-descriptiveness or performed a vowel-counting control task. | self > non-self-referential attention control | Yes |
| McAdams and Krawczyk (2014) | anorexia nervosa | 18 | 18 | social adjectives | Appraise self, friend, and reflected perspectives in social and physical identity tasks. | self > friend; self > reflected-self | Yes |
| Komulainen et al. (2018) | major depressive disorder | 15 | 15 | Positive, negative, and neutral adjectives | Participants categorized self-referential emotional adjectives, with neutral words serving as controls. | self-referential > neutral | Yes |
| Dixon et al. (2022) | social anxiety disorder | 34 | 97 | Positive/negative trait words | Participants made yes/no judgments on balanced positive/negative trait adjectives for self-descriptiveness or letter case. | positive self > positive case judgment; negative self > negative case judgment | No |
| Li et al. (2017) | major depressive disorder | 21 | 19 | Positive/negative personality trait adjectives | Participants judged whether positive or negative personality adjectives described themselves (self-referential condition) or were socially desirable (semantic/other-referential condition). | positive self > positive case judgment; negative self > negative case judgment | No |
| Hur et al. (2021) | social anxiety disorder | 22 | 25 | self-referential social evaluation | Participants performed self-referential processing in a virtual social setting. | positive > neutral self-relevance; negative > neutral self-relevance | No |
| Beeney et al. (2016) | personality disorder | 21 | 17 | trait words | Participants were asked to identify a friend with whom they were close. Evaluation targets were arranged in a 2×2 factorial design with the following conditions: (a) first-person, self (Are you kind?), (b) first-person, other (Is Julie kind?), (c) third-person, self (According to Julie, are you kind?), and (d) third-person, other (According to Julie, is she kind?). | self > close other; personality > physical traits; referent | No |
| Bluhm et al. (2012) | post-traumatic stress disorder | 15 | 20 | self-characteristic and general-fact sentences | Participants responded yes/no to self-related, other-related and general-fact auditory statements. | self-characteristics > general facts; self-characteristics > other-characteristics | No |
| Holt et al. (2011) | schizophrenia | 17 | 18 | self/other trait judgments | Participants completed self-reflection, affect labeling, mother-referential reflection and letter-case perceptual judgments on matched trait adjectives in an 18-s block fMRI task. | self-reflection > affective appraisal; self-reflection > other-reflection | No |
| Kennedy and Courchesne (2008) | autism spectrum disorder | 12 | 13 | self/other mentalizing judgments | Participants judged true or false for internal trait and external behavioral statements about themselves or their mother, completed math baseline trials and rest viewing in this block fMRI paradigm. | self > other; self > letter-case control; other > letter-case control | No |
| Komeda et al. (2015) | autism spectrum disorder | 15 | 15 | trait judgments | Participants completed self-agreement or self-character similarity judgments on sentences describing gender-matched autistic or non-autistic fictional characters. | self-related > other-related narrative/mental-state conditions | No |
| Lythe et al. (2022) | bipolar disorder | 39 | 67 | self-referential words | Participants rated unpleasantness for moral violation sentences featuring self or their best friend as the agent during fMRI, with a passive asterisk baseline; post-scan ratings and emotion choices were used to select high-unpleasantness trials. | self-blame > other-blame; self-blame > visual baseline; other-blame > visual baseline | No |
| Shad and Keshavan (2015) | schizophrenia | 15 | 17 | trait statements | The self-assessment task used self-referential and other-referential cue epochs with visual self- and other-directed sentences, requiring participants to judge whether the presented remarks refer to themselves or other people. | self-directed > other-directed statements | No |
| Belleau et al. (2015) | major depressive disorder | 16 | 16 | self-referential adjectives | Participants completed self-focused abstract rumination-style and external-focused visual-imagery thought-induction tasks inside the scanner, with task order counterbalanced. | self-focused > externally focused thought induction | No |

For each eligible task contrast, we extracted the reported anatomical labels, peak coordinates, coordinate space, statistical thresholds and correction procedures. Direct group differences (psychiatric patient > healthy control and psychiatric patient < healthy control) were coded as higher or lower activation in psychiatric patients relative to healthy controls. Neuroimaging acquisition and analysis characteristics are summarized in Supplementary Table S3.

### 2.3 Systematic summary of included studies

To provide a comprehensive overview of differences in brain activation between psychiatric patients and healthy controls during self-reflection, we systematically summarised key characteristics and reported findings of the included studies.

We manually checked all included articles and coded two types of information: (1) whether the study reported altered patient-control differences for self versus other or self versus control task contrasts for a given psychiatric diagnosis, and (2) whether the reported task-contrast estimates were higher or lower in psychiatric patients relative to healthy controls. Findings were organized by diagnostic category and interpreted alongside the exact task contrast. Findings concerning group differences across conditions, connectivity, clinical associations, or treatment-related changes were distinguished from direct patient-control task-contrast differences.

### 2.4 Activation likelihood estimation (ALE) based meta-analysis

To identify brain regions exhibiting convergent above-chance spatial convergence of patient-control differences across diagnostic disorders, we carried out an activation likelihood estimation (ALE) meta-analysis (Acar et al., 2018; Eickhoff et al., 2009, 2012, 2016; Turkeltaub et al., 2012).

The ALE algorithm evaluates whether reported activation foci converge across experiments at a rate greater than expected by chance within a gray-matter mask. In ALE, each reported peak is modeled as the center of a three-dimensional Gaussian probability distribution, with kernel width scaled by sample size to account for between-subject and between-template variability (Eickhoff et al., 2009). For each experiment, a modeled activation (MA) map is generated by assigning the highest probability from any focus to each voxel. The union of MA maps across experiments yields a whole-brain ALE map representing convergence of activation likelihoods. This ALE map is compared against a null distribution generated under the assumption of random spatial association among experiments, and significance is assessed using random-effects inference (Eickhoff et al., 2016b; Turkeltaub et al., 2012). To control for multiple comparisons, we applied cluster-level family-wise error (FWE) correction, which balances Type I error control and statistical power.

All analyses were implemented in NiMARE (Salo et al., 2023) (v0.20.0) in MNI152 space at 2 × 2 × 2 mm resolution (Fonov et al., 2011). Seven of these experiments reported foci in Talairach space and were converted to MNI space using the inverse Lancaster information (Lancaster et al., 2007), as implemented in GingerALE (v3.0.2). We applied a voxel-level threshold of *p* < 0.001 (uncorrected) and a cluster-level threshold of *p* < 0.05 (FWE-corrected). The latter was determined by constructing an empirical distribution of maximum cluster sizes from 10,000 iterations in which random peak locations were drawn from a gray matter template.

For each task-contrast category, we conducted ALE analyses pooling coordinates from both directions of patient–control differences and separately analyzing higher and lower activation in patients relative to healthy controls. These analyses assessed spatial convergence irrespective of direction and within each direction. Exploratory ALE contrast analyses compared self-versus-control with self-versus-other datasets separately for each patient–control direction. Full details are given in Supplementary Section S10.

We followed the recommendations regarding statistical power in ALE analysis (Eickhoff et al., 2016; Frahm et al., 2022) and reported experiment counts and the heuristic sample-size-adjusted dataset counts described by Radua et al. (2026) (see Supplementary Section S6). To characterize the contribution of individual experiments to significant clusters, we used the FocusCounter function in NiMARE (v0.20.0) to identify experiments with at least one reported peak within each cluster (see Supplementary Table S6). We additionally conducted exploratory meta-analytic functional decoding of the unthresholded ALE *z*-maps to describe their spatial associations with psychological terms (see Supplementary Section S12 for methodological details).

Volumetric visualizations in MNI152 space were performed using Nilearn (Abraham et al., 2014) (v0.9.2), and anatomical labels were assigned using AtlasReader (Notter et al., 2019) (v0.1.2) with the Automated Anatomical Labelling Atlas 3 (AAL3) (Rolls et al., 2020). Surface projections were generated in Connectome Workbench (Glasser et al., 2016). The full pipeline, including analysis code and visualization scripts, is openly available on GitHub (https://github.com/Chuan-Peng-Lab/Self_Psych_Meta.git).

## 3 Results

### 3.1 Overview of included studies

The systematic review comprised 36 studies, of which 25 were eligible for ALE analyses and 11 were included in the systematic review only (Table 1). The eight diagnostic categories were schizophrenia disorders (n = 16 experiments), depressive disorders (n = 7), autism spectrum disorder (n = 4), anxiety disorders (n = 4), bipolar disorders (n = 2), feeding or eating disorders (n = 2), trauma- and stressor-related disorders (n = 1), and personality disorders (n = 1). Note that Zhang et al. (2015) included schizophrenia and bipolar groups and is counted in both categories but only once in the total of 36 studies.

Studies differed in clinical populations, stimuli, comparison conditions, and neuroimaging analysis procedures. Table 1 describes the study-level tasks and samples, Table S3 describes acquisition and analysis characteristics, and Table S4 summarizes reported regional directions.

The six ALE analyses synthesized patient–control differences separately by task-contrast category and effect direction, considering both pooled and direction-specific results. Table 2 reports their dataset composition.

**Table 2.** Composition of the datasets included in the single ALE analyses.

| Dataset | Experiments | Sample size | Sample-size-adjusted dataset count | Foci |
|---|---|---|---|---|
| Patients<sub>[self>control]</sub> < healthy controls<sub>[self>control]</sub> | 9 | 363 | 4.67 | 18 |
| Patients<sub>[self>other]</sub> < healthy controls<sub>[self>other]</sub> | 8 | 331 | 4.21 | 18 |
| Patients<sub>[self>control]</sub> > healthy controls<sub>[self>control]</sub> | 3 | 110 | 1.45 | 5 |
| Patients<sub>[self>other]</sub> > healthy controls<sub>[self>other]</sub> | 8 | 225 | 2.54 | 23 |
| Both directions pooled; [self>control] | 12 | 473 | 6.12 | 23 |
| Both directions pooled; [self>other] | 13 | 491 | 6.17 | 41 |

***Note.*** Sample size denotes the total number of participants across the psychiatric patient and healthy control groups in the included experiments. Sample-size-adjusted dataset count was calculated using the heuristic proposed by Radua et al. (2026) (see Supplementary Section S6).

### 3.2 Results of systematic review

Across 36 studies, a wide range of aberrant brain activations during self-reflection was reported in the eight broad categories of psychiatric disorders. The main findings are described below.

1. Schizophrenia Spectrum Disorders. In 11 of the 16 included studies, within-group analyses reported greater neural responses during the self versus other condition in cortical midline and lateral regions, including the posterior cingulate cortex (PCC), medial prefrontal cortex (mPFC), superior frontal gyrus (SFG), angular gyrus, precuneus, and middle frontal gyrus (MFG) (Bedford et al., 2012; Blackwood et al., 2004; Furuichi et al., 2019; Holt et al., 2011; Jimenez et al., 2018; Liu et al., 2014; Pauly et al., 2014; Shad et al., 2012; Tan et al., 2015; van der Meer et al., 2013; Zhang et al., 2015). Patients-control group comparisons also showed differences in both directions. Higher self versus other contrast estimates in patients were reported in the supplementary motor areas (SMA), anterior cingulate cortex (ACC), and frontal, parietal and posterior cingulate gyrus, whereas lower estimates were reported in superior frontal gyrus, temporal gyrus and medial prefrontal cortex, including the ventromedial prefrontal cortex (Bedford et al., 2012; Blackwood et al., 2004; Fuentes-Claramonte et al., 2020; Furuichi et al., 2019; Holt et al., 2011; Liu et al., 2014; Pankow et al., 2016; Pauly et al., 2014; Shad et al., 2012; Tan et al., 2015). Makowski et al. (2016) additionally found lower self versus other contrast estimates in patients in right parietal regions, including the angular gyrus and superior and inferior parietal regions, under the high social reward condition. For self versus control, Murphy et al. (2010) reported lower activation in patients in the precuneus, angular gyrus, and lingual gyrus relative to semantic positivity judgments. Some studies additionally examined connectivity or associations with insight, which address different outcomes from the direct group contrasts (Holt et al., 2011; Shad & Keshavan, 2015b; van der Meer et al., 2013b).
2. Depressive and anxiety disorders. For depressive disorders, Lemogne et al. (2009) reported higher activation in the middle frontal gyrus in patients for the self versus control contrast. Across three studies, patients showed higher activation relative to healthy controls in the dorsomedial prefrontal cortex (dmPFC), inferior frontal gyrus (IFG), anterior subgenual cingulate cortex (sgACC), superior temporal gyrus (STG), and medial superior frontal cortex (mSFG), with no regions showing lower activation (Lemogne et al., 2009; Lythe et al., 2022; Sarsam et al., 2013). No regions of lower activation were reported in these three studies. However, Li et al. (2017) found higher activation in the central medial prefrontal cortex (mPFC) and lower activation in the dorsal mPFC in patients. Belleau et al. (2015) reported no significant group differences. Emotional valence and treatment were additional dimensions in studies by Yoshimura et al. (2010) and Komulainen et al. (2018). For anxiety disorders, three studies found that the amygdala, medial prefrontal cortex (MPFC), anterior cingulate cortex (ACC), lateral middle frontal cortex, and medial frontal gyrus were more activated in the self-than in the other-referential condition in the patient group (Blair et al., 2008; Blair et al., 2011; Pujol et al., 2013). Also, two studies reported higher activation in patients in the middle occipital gyrus, ventral MPFC, medial temporal cortex, and medial frontal cortex (Hur et al., 2021; Pujol et al., 2013), and one study observed lower activation in patients in the prefrontal cortex and medial frontal cortex (Pujol et al., 2013). Hur et al. (2021) further reported that patients showed higher activation in the bilateral inferior frontal gyrus and left middle temporal gyrus for positive versus neutral self-referential words, whereas they showed higher left insular activation and lower right middle frontal gyrus activation for negative versus neutral self-referential words. Notably, Dixon et al. (2022) found no significant group differences.
3. Neurodevelopmental disorders. Two studies found not differences in neural activation between groups for the self versus other condition (Kennedy & Courchesne, 2008; Komeda et al., 2015). Conversely, two studies reported lower activation in patients than in healthy controls in the middle cingulate cortex (MCC) (Hashimoto et al., 2017; Lombardo et al., 2010b). However, Hashimoto et al. (2017) examined functional connectivity using self and mother as referential targets from first- and third-person perspectives.
4. The other disorders. For bipolar and related disorders, one study found no significant group differences for the self versus control contrast (Zhang et al., 2015), but Porta-Casteràs et al. (2023) found that bipolar patients showed greater dorsomedial and dorsolateral prefrontal cortex activation than healthy controls during the self versus control condition. For feeding and eating disorders, McAdams & Krawczyk (2014) found reduced precuneus and anterior cingulate activation during self-related identity appraisal in patients. McAdams et al. (2016) further reported greater mPFC and dorsal ACC responses for the self versus friend contrast in currently ill women relative to healthy controls. For trauma- and stressor-related disorders, Bluhm et al. (2012) identified significantly higher activation in the dorsomedial prefrontal cortex (DMPFC) and precuneus during the self versus other condition. They also reported lower activation in the ventromedial prefrontal cortex (VMPFC) when comparing patients with healthy controls (Bluhm et al., 2012). Finally, for personality disorders, Beeney et al. (2016) found no brain regions showing higher activation for the self versus other condition among patients, but observed lower activation in visual, sensory, motor, and mirror-neuron–related regions.

Across diagnostic categories, reported patient–control task-contrast differences were distributed across cortical midline and lateral association regions. Medial frontal and prefrontal regions were reported for both self versus other and self versus control comparisons and for both higher and lower task-contrast estimates in patients. Cingulate and precuneus findings were more frequently reported as lower patient task-contrast estimates, although reports in the opposite direction were also present in the posterior and middle cingulate cortices. Lateral frontal and parietal regions were represented in both comparison categories, whereas temporal, limbic, sensorimotor, and visual regions were reported less consistently (see Figure 2).

**Figure 2.**
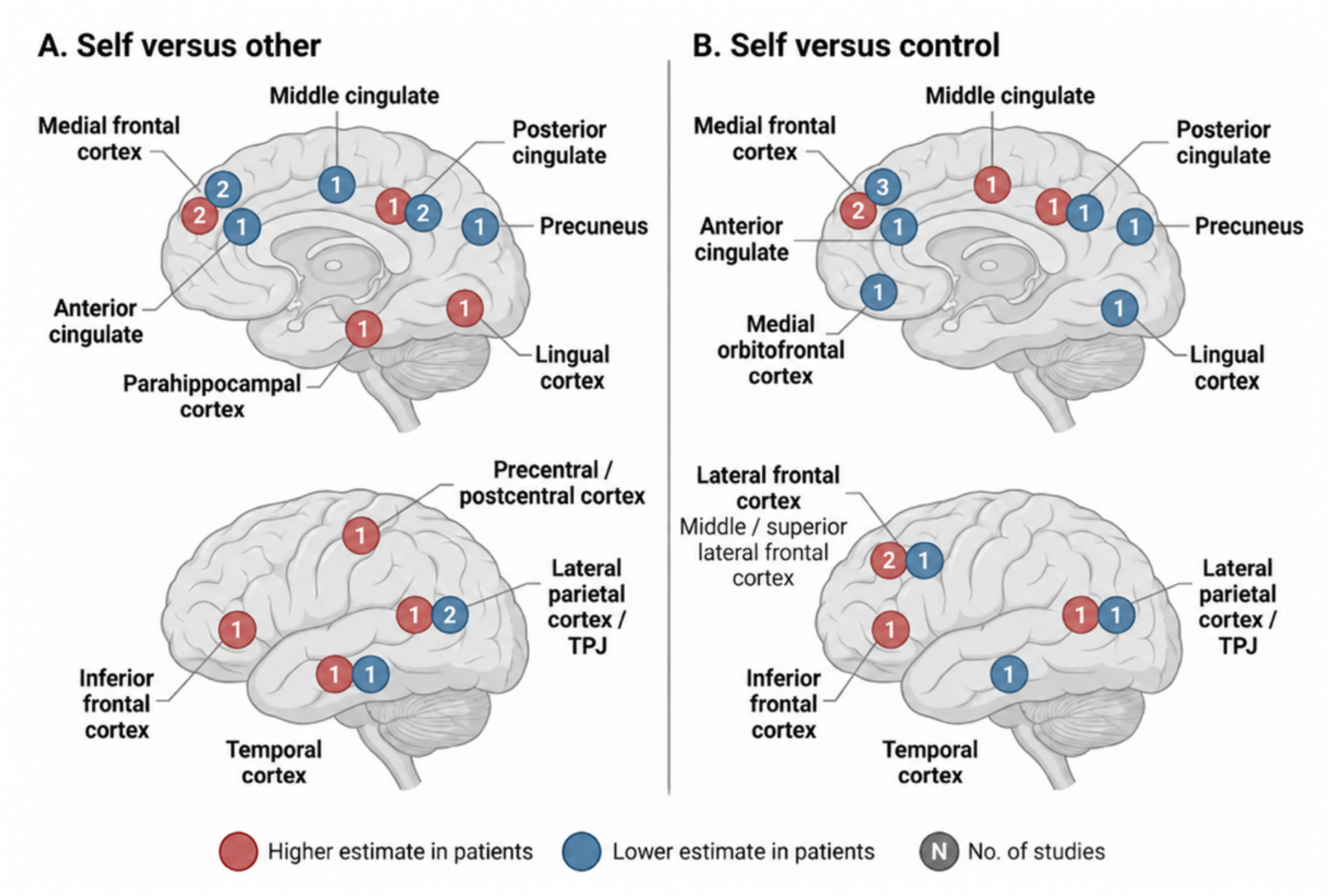
Main regional distribution and direction of reported patient–control differences. ***Note.*** Red circles indicate higher activation in patients than in healthy controls, whereas blue circles indicate lower activation. The source studies, exact contrasts, and analysis scopes are listed in Supplementary Tables S3–S5. This figure was created with BioRender.com (https://www.biorender.com/).

### 3.3 Results of ALE analysis

The overall ALE identified a cluster centered in the right anterior cingulate cortex (ACC) for self-versus-other contrast and a cluster in the left middle frontal gyrus (MFG) for the self-versus-control contrast (see Table 2). FocusCounter analyses revealed that 3 of 13 experiments (23.1%) had peaks inside the right ACC, all from schizophrenia samples. In addition, 3 of 12 experiments (25.0%) had peaks inside left MFG, two of these involved schizophrenia and one involved depression.

**Table 2.** Significant clusters in the ALE analyses.

| Peak region | Size (mm<sup>3</sup>) | Peak z | Peak MNI coordinate |  |  | Cluster | Contributing experiments(*n*/*N*) |
|---|---|---|---|---|---|---|---|
|  |  |  | x | y | z |  |  |
| **Single ALE: higher activation and lower activation contrast; self vs other** |  |  |  |  |  |  |  |
| R anterior cingulate cortex | 720 | 5.04 | 8 | 38 | 0 | R anterior cingulate cortex (92.22%); R medial orbitofrontal cortex (5.56%) | 3/13(23.1%) |
| **Single ALE: higher activation and lower activation contrast; self vs control** |  |  |  |  |  |  |  |
| L middle frontal gyrus | 768 | 4.20 | -44 | 16 | 40 | L middle frontal gyrus (78.12%); L precentral gyrus (18.75%) | 3/12(25.0%） |
| **Single ALE: psychiatric patient < healthy control; self vs other** |  |  |  |  |  |  |  |
| L anterior cingulate cortex | 536 | 4.43 | -6 | 46 | -4 | L medial orbitofrontal cortex (53.73%); L anterior cingulate cortex (46.27%) | 2/8(25.0%） |
| **Single ALE: psychiatric patient < healthy control; self vs control** |  |  |  |  |  |  |  |
| L middle frontal gyrus | 488 | 4.26 | -44 | 16 | 40 | L middle frontal gyrus (81.97%); L precentral gyrus (13.11%) | 2/9(22.2%) |

For results from the patients < healthy control comparison, self-versus-other yielded a cluster centered in the left ACC/orbitofrontal, and self-versus-control yielded a cluster centered in the left MFG (Table 2 and Figure 3). No significant clusters were identified for the patients > healthy control comparison. FocusCounter analyses revealed that the left ACC/orbitofrontal region contained peaks from 2 of 8 experiments (25.0%), involving autism and schizophrenia, whereas the left MFG contained peaks from 2 of 9 experiments (22.2%), both involving schizophrenia (see Supplementary Table S6).

**Figure 3.**
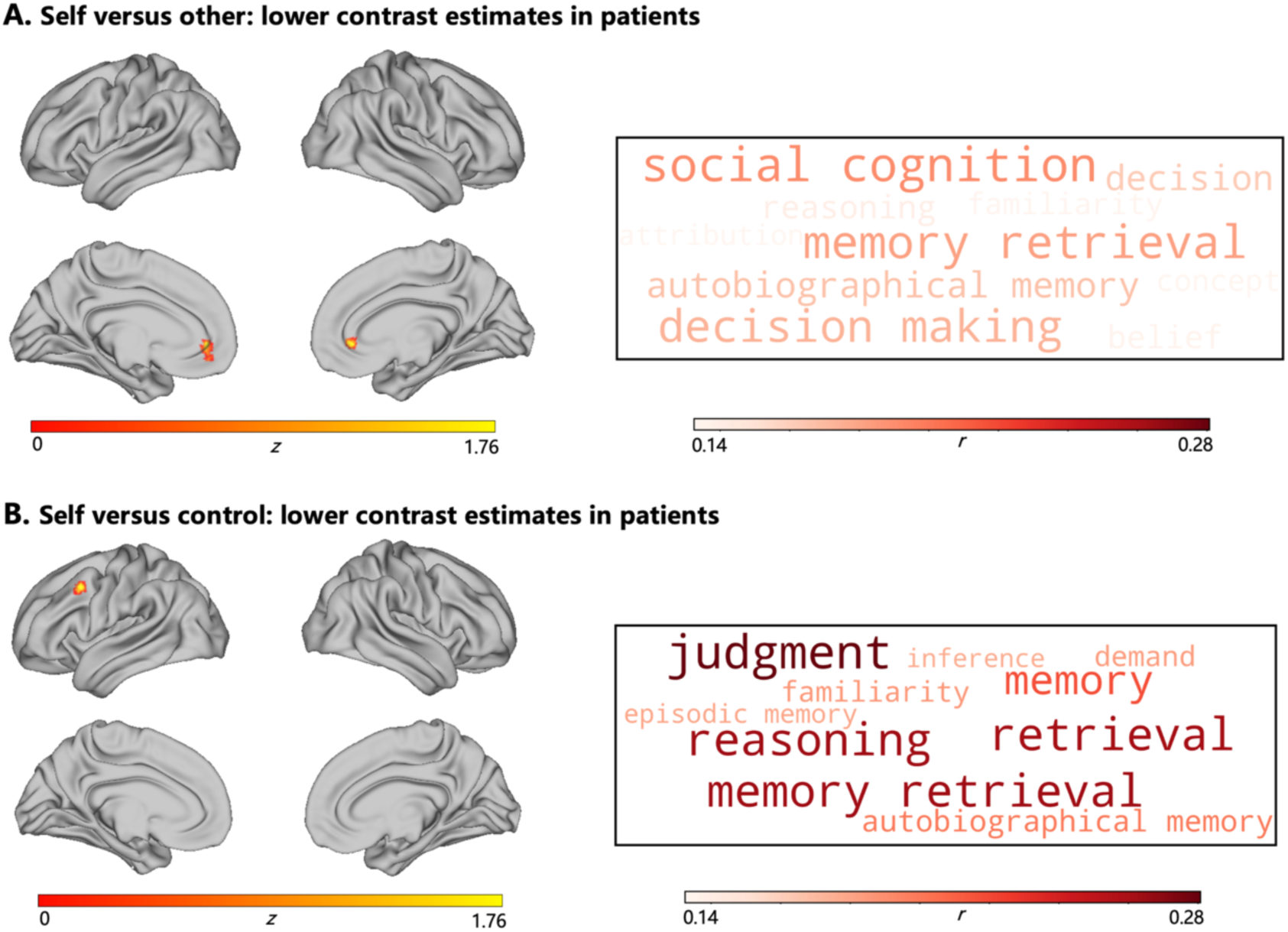
Spatial convergence of lower patient–control task-contrast estimates and exploratory functional decoding. ***Note.*** Brain maps show significant ALE clusters at a cluster-level FWE-corrected threshold of *p* < .05, with a cluster-defining voxel-level threshold of *p* < .001 (uncorrected). Word clouds display the 10 terms with the highest positive Pearson correlations with the corresponding unthresholded ALE *z* maps.

Functional decoding based on these two significant clusters revealed that the left ACC/orbitofrontal cortex was associated with “social cognition”, “memory retrieval” and “decision making”. The left MFG was associated with “judgment”, “reasoning” and “memory retrieval” (Figure 3). Exploratory contrast analyses found no significant differences in spatial convergence between the self-versus-control and self-versus-other datasets, either for patient < control contrast effects or for patient > control contrast effects, in either subtraction direction.

## 4 Discussion

To our knowledge, this is the first systematic review and meta-analysis to assess altered neural responses during self-reflection across a broad range of psychiatric disorders. Through the systematic review, we found that altered neural responses during self-reflection were reported across multiple psychiatric disorders, echoing the widely observed self-related difficulties across psychiatric disorders in the literature (Philippi & Koenigs, 2014; Zhao et al., 2013). The ALE analyses also identified two brain regions, the ACC and MFG, across these psychiatric disorders during self-reflection. However, these findings cannot provide solid evidence for the notion that neural responses during self-reflection are transdiagnostic features. Thus, our findings identify important issues in the field rather than providing answers to these questions.

First, we observed that studies reporting altered neural responses during self-reflection covered many psychiatric disorders, confirming the broad interest in self-reflection across psychiatric disorders. Previous studies have reported differences in self-related mental processes between individuals with psychiatric disorders and control groups. For example, depression is often associated with negative self-evaluation and self-critical rumination, whereas social anxiety commonly involves heightened self-focused attention and concern about how one is perceived by others (Blair et al., 2008; Derry & Kuiper, 1981; Nolen-Hoeksema et al., 2008; Tamura et al., 2025). In schizophrenia, difficulties in evaluating one’s own experiences and characteristics have been discussed in relation to impaired self-awareness, with both heightened and diminished self-reflection reported (Furuichi et al., 2019; van der Meer et al., 2013). These varied clinical manifestations may have motivated researchers to explore the relationship between self-reflection and psychiatric disorders. As the self-referential encoding task (Rogers et al., 1977) is widely used to examine the neural basis of self-reflection (see Hu et al., 2016; Sun et al., 2023), it is unsurprising that this task has also been adopted to examine differences in neural responses during self-reflection across psychiatric disorders.

Complementing the qualitative findings, our ALE analyses identified spatial convergence in the ACC for self versus other contrasts. This brain region appeared both in the ALE analysis that pooled all altered between-group differences and in the ALE analysis that specifically focused on lower activation during self-reflection in psychiatric patients than in healthy controls. The central role of the ACC in self-reflection and related processes has been widely reported in the literature (Araujo et al., 2013; Hu et al., 2016; Northoff et al., 2006; Qin et al., 2020). For example, Hu et al., (2016) have reported the ACC in self-reflection, as compared to self-face processing, among healthy participants. Qin et al (2020) also found that ACC supports the mental-self process, but is less involved interoceptive-processing or exteroceptive-processing. Our results suggest that the ACC might be an important hub for task-induced self-reflection and that its activation pattern may differ between psychiatric patients and healthy controls. This finding is consistent with the RDoC framework, under which Pintos Lobo et al. (2023) found the ACC to be one of the regions preferentially associated with the broader construct of perception and understanding of the self. Taken together, these findings suggest that the ACC may provide a candidate region for further investigation of altered self-reflection in psychiatric disorders.

On the other hand, our study identified equally prominent questions in the field. The most obvious finding is the uneven distribution of studies across diagnoses. We found that schizophrenia accounted for the largest share of studies, followed by depressive disorders, anxiety disorders, and autism spectrum disorder. In schizophrenia, disturbances of self-experience have been salient in clinical descriptions and have subsequently become the focus of systematic research programmes (Ebisch & Aleman, 2016; J. Liu et al., 2014; Lysaker et al., 2013; Nordgaard & Parnas, 2014). In depression and social anxiety, rumination and self-focused cognition have likewise been incorporated into influential models of symptom development and maintenance (Nolen-Hoeksema et al., 2008; Norton & Abbott, 2016). Previous systematic reviews and meta-analyses have also focused on these diagnostic groups (Potvin et al., 2019; Sabbah & Northoff, 2024, 2025; Zhou et al., 2026). Other disorders, however, have received far less attention. For example, although negative self-evaluation and low self-esteem are involved in the development and maintenance of anorexia nervosa, most studies exploring self-disturbance in this condition focus primarily on self-image (Gu et al., 2024; Kästner et al., 2019). This uneven distribution of studies means that we still lack enough data for transdiagnostic claims. Given the importance of self in mental processes and mental health (Northoff et al., 2026), our findings call for more studies on these less well-represented diagnoses.

Another issue we identified concerns how self-reflection is operationalized. The self is a multidimensional construct, and its conceptual boundaries differ across research traditions (Hards et al., 2024; Klein, 2012; Qin et al., 2020), and the terminology is not often clear (van der Meer et al., 2010; Knyazev et al., 2021). Previous meta-analyses have often combined studies using tasks that probe different self-related processes (Potvin et al., 2019; Sabbah & Northoff, 2024, 2025; Zhou et al., 2026). These studies indeed provide valuable knowledge about self-related disturbances, but they also make it difficult to distinguish which aspects of self-related processes are altered across disorders. Such distinctions may also help identify more specific targets for intervention (Kiluk, 2018). We focused on self-reflection as a step forward. Yet variation in task implementation emerged as a new challenge (Sun et al., 2023). In our ALE results, spatial convergence in the MFG was identified for self-versus-control contrasts. However, control conditions varied across studies, for example, making true or false judgements about general-knowledge statements (Fuentes-Claramonte et al., 2020) or judging whether neutral trait adjectives contained five or more vowels (Porta-Casteràs et al., 2023). Thus, the self-versus-control contrasts may capture differences in semantic processing, evaluative demands, or response selection, in addition to self-related processing. This limits the specificity with which the MFG finding can be attributed to self-reflection. Together, our study highlights inconsistencies in the operationalization of self-related processed and calls for clearer definitions and more explicit links between theoretical constructs and experimental measures.

When interpreting our findings, the following limitations should be noted. First, the number of experiments in each ALE analysis, defined by task contrast and effect direction, fell below the recommended number (Eickhoff et al., 2016; Müller et al., 2018), limiting the statistical power of our ALE analyses and potentially increasing the probability of false negatives or false positives. Second, ALE assesses spatial convergence rather than individual effect sizes. Thus, the lack of any significant cluster for the directional ALE analysis should be interpreted with caution.

Based on these findings, we suggest that future studies should define the exact aspects of self-related processing being examined and use comparable tasks across diagnostic groups, with clearly specified comparison targets, stimulus valence, and task demands (Sun et al., 2023). Reporting responses to both the self and comparison conditions would help clarify the source of group differences. Future studies should also systematically collect fMRI data from underrepresented clinical populations. Multicenter, multidimensional collaborations with open data could increase sample diversity, improve diagnostic representation, and enhance the transparency, generalizability, and robustness of cross-diagnostic findings (Alexander et al., 2017; Chopra et al., 2025; Qi et al., 2025).

