## Supplementary Materials for "Do altered neural responses during self-reflection converge across psychiatric disorders? A systematic review and meta-analysis of neuroimaging studies"

MS Shanshan Zhu[1], PhD Wen-Jing Yan[2], PhD Hu Chuan-Peng[1, 3]
1 School of Psychology, Nanjing Normal University, 210024 Nanjing, China;
2 School of Mental Health, Wenzhou Medical University, Wenzhou, China;
3 The Fourth People's Hospital of Yancheng, Affiliated Yancheng Hospital, Nanjing Normal University, Institute of Brain Science and Mental Health, Yancheng, China

**Author Note**

Shanshan Zhu https://orcid.org/0009-0007-6612-6190

Wen-Jing Yan https://orcid.org/0000-0001-6413-2414

Hu Chuan-Peng https://orcid.org/0000-0002-7503-5131

**S7. Supplementary results: fMRI acquisition and analysis characteristics of the 36 included studies ..................................... ............................................................ 10**

### S1. Supplementary methods: Sources and search terms.

Table S1 summarizes the literature search strategy used in the present review, including the databases searched and the keyword combinations applied to identify neuroimaging studies on self-reflection in psychiatric patients and healthy controls.

**Table S1.** Sources and search terms.

| Search | Databases | Keywords |
|---|---|---|
| Human adults with psychiatric disorders and healthy controls | PubMed, Web of Science, NeuroSynth, BrainMap, EMBASE, SCOPUS | ("self-referen*" OR "self-assessment" OR "self-perception" OR "self-evaluation" OR "self-confidence" OR "self-appraisal" OR "self-awareness" OR "self-concept" OR "self-representation" OR "self-esteem" OR "self-recognition" OR "self-criticism" OR "self-reflection" OR "self-related processes" OR "self-identity") AND ("fMRI" OR "functional magnetic resonance imaging" OR "PET" OR "positron emission tomography"). |

### S2. Supplementary methods: Coding procedure and inter-rater reliability assessment.

The dataset was independently coded by two trained researchers to minimize subjective bias and enhance methodological rigor. Following completion of the independent coding process, the coders cross-checked all entries and assigned a binary indicator (0 = non-identical; 1 = identical) to reflect agreement on each item. Inter-rater reliability was subsequently quantified based on these agreement scores. The source code used for agreement scoring and reliability estimation is provided in the online supplementary materials to ensure transparency and reproducibility.

As a meta-research-based dataset, the core of its quality resides in the accuracy and consistency of the coding process. In this study, we employed the $AC_1$ coefficient proposed by (Gwet, 2008) to quantify inter-rater reliability. The $AC_1$ coefficient is considered more robust than the traditional Cohen's Kappa coefficient (Wongpakaran et al., 2013), particularly in handling skewed distributions of categories. The calculation formula is as follows:

$$AC_1 = \frac{p_a - p_{e\gamma}}{1 - p_{e\gamma}} \quad (1)$$

Where $p_a$ represents the overall probability of agreement (including both chance and non-chance agreement), and $p_{e\gamma}$ denotes the probability of chance agreement. The $AC_1$ coefficient was calculated using the *irrCAC* package in R v4.5.2.

### S3. Supplementary results: Deviations from the preregistration.。

**Table S2.** Preregistration Deviations Table.

| **Deviations** | | | | | |
|---|---|---|---|---|---|
| # | Details | | Original Wording | Deviation Description | Reader Impact |
| 1 | Type | Data Preparation | *If a patient is formally diagnosed with a mental disorder by DSM or ICD criteria, it is considered that the patient has a mental illness.* | In the final version, we operationalised this definition by requiring studies to include participants diagnosed using standardised diagnostic instruments or criteria (e.g., DSM-IV-TR, DSM-IV, ICD-10, MINI-Plus, or ADOS). | This deviation is unlikely to affect readers' interpretations of the findings, as it represents a clarification and operational refinement of the original definition rather than a substantive change to the inclusion criteria. In addition, for autism spectrum disorder, ADOS is considered a gold-standard diagnostic instrument (Lord et al. 2008). |
| | Reason | Other (Please Explain) | | | |
| | Timing | During data collection | | | |
| | Reason | Other (Please Explain) | | | |
| | Timing | Before data collection | | | |
| 2 | Type | Analysis | *Additional outcome(s): We conducted* | We assessed the robustness of our | This deviation is unlikely to affect the |

| | | | | | |
|---|---|---|---|---|---|
| | Reason | Other (Please Explain) | *leave-one-out analysis and Fail-safe N analysis to test the robustness of our results and the potential effect of publication bias.* | meta-analytic results and the potential impact of publication bias using the Python package NiMARE, specifically employing FocusCounter and fail-safe *N* analysis. | interpretation of our results, as the diagnostic analyses serve the same purpose of assessing result stability and potential publication bias. |
| | Timing | After data access | | | |

**S4. Supplementary results: The geographical distribution of psychiatric patients.**

Based on the geographic distribution of the included samples, the largest number of participants were recruited from the United States, which contributed the highest patient sample overall. In total, the dataset comprised 818 psychiatric patients drawn from 12 different countries.

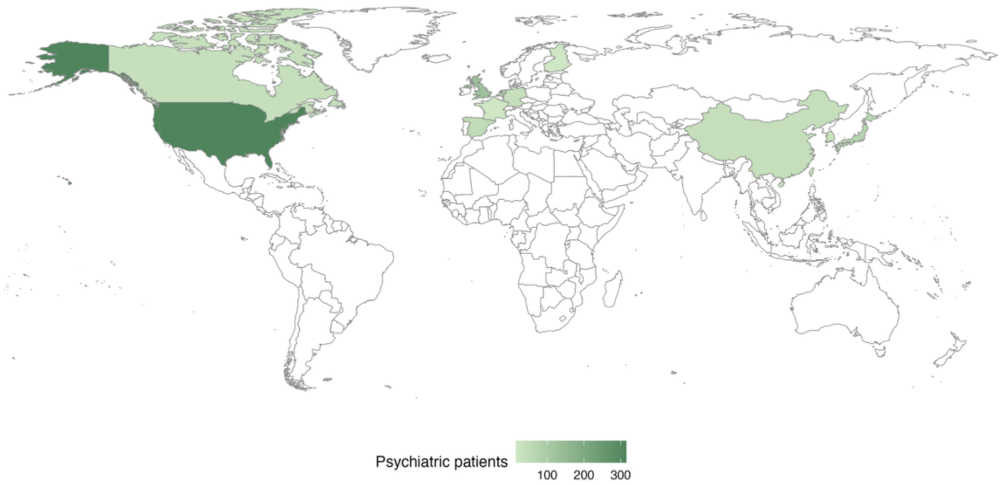

**Figure S1.** The geographical distribution of psychiatric patients and the sample sizes for the studies included.

### S5. Supplementary results: Distribution of study articles and psychiatric patient sample sizes across psychiatric disorder types.

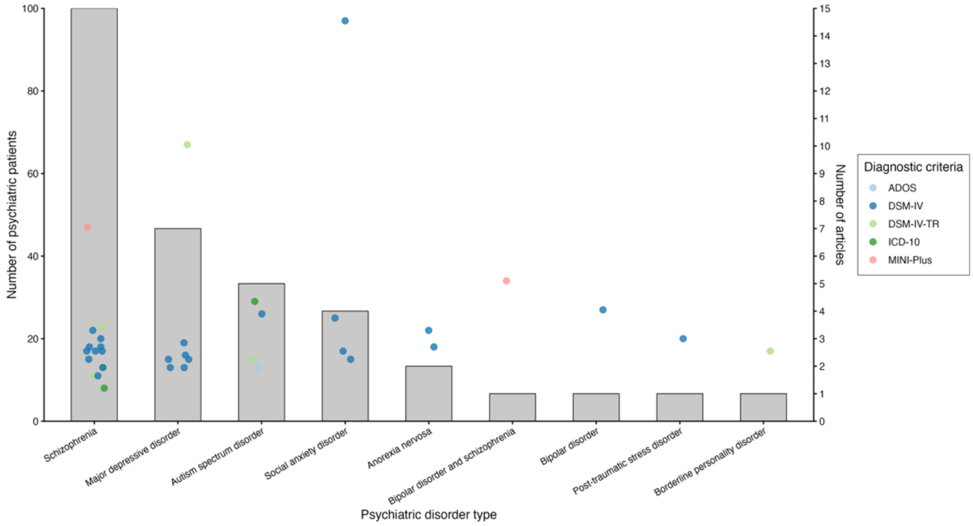

**Figure S2.** Distribution of study articles and psychiatric patient sample sizes across psychiatric disorder types.

***Note.*** MINI-Plus: Mini International Neuropsychiatric Interview-Plus; ADOS: Autism Diagnostic Observation Schedule.

### S6. Supplementary methods: Adjustment for the sample size of included experiments when counting the number of datasets.

A standard ALE meta-analysis has been recommended to include approximately 17 independent peak-coordinate datasets to achieve adequate statistical power (Eickhoff et al., 2016; Frahm et al., 2022, 2023). Because our directional ALE analyses included fewer than 17 independent datasets, and the included datasets varied considerably in sample size, we additionally applied the heuristic sample-size adjustment proposed by Radua et al. (2026), adapting the R code provided in their Supplementary Material.

Under this approach, each independent dataset is assigned a sample-size-adjusted count equal to its estimated statistical power divided by a conventional reference power of 80%. Following Radua et al. (2026), power was calculated for a one-sided independent-samples t-test assuming a very large effect (Cohen's $d$ = 1) and a significance level of $\alpha$ = .001:

$$\text{Adjusted dataset count}_i = \frac{Power_i}{0.80}$$

Consequently, datasets with smaller samples contribute less than one to the adjusted count, a dataset with 80% power contributes one, and the contribution of an individual dataset cannot exceed 1.25 studies. Because the original calculation assumed equal group sizes, we adapted it to use the actual, and sometimes unequal, numbers of psychiatric and healthy-control participants in each dataset (see our code available on GitHub). This procedure provides a heuristic sample-size-adjusted count of the available datasets rather than a formal estimate of the overall statistical power of the ALE analysis.

### S7. Supplementary results: fMRI acquisition and analysis characteristics of the 36 included studies.

**Table S3.** fMRI acquisition and analysis characteristics of the 36 included studies.

| Study | Disorder | Preprocessing | Multiple-comparison correction | Cluster threshold | Software | Tesla |
|---|---|---|---|---|---|---|
| Study | Disorder | Discard first 10 volumes; motion correction (MCFLIRT); functional–structural coregistration; MNI normalization; 5-mm FWHM smoothing | Gaussian random-field cluster correction | $Z > 2.3$; cluster-corrected $p < .05$ | FSL5.0 | 3T Philips Achieva |
| Furuichi et al. (2019) | Schizophrenia | Discard first 8 images; realignment; T1 coregistration; MNI normalization; 8-mm FWHM smoothing; 128-s high-pass filter | Cluster-level FWE | Voxelwise $p < .001$ uncorrected; cluster FWE $p < .05$ | SPM8 | 1.5T Siemens Magnetom Vision |
| Fuentes-Claramonte et al. (2020) | schizophrenia | Motion-artifact correction; 8.08-mm FWHM smoothing; spatial normalization to standard space; permutation-based analysis | Permutation-based cluster inference | Voxelwise $p < .05$ uncorrected | XBAM | 1.5T GE Signa |

| | | | | | | |
|---|---|---|---|---|---|---|
| Furuichi et al. (2019) | schizophrenia | Realignment; spatial normalization; 10-mm FWHM smoothing | Whole-brain correction; SVC for group contrasts | Whole-brain $p < .05$ corrected; group contrasts SVC $p < .01$ corrected | SPM99 | 1.5T GE NV/i Signa LX Horizon |
| Bedford et al. (2012) | schizophrenia | Discard first 4 volumes; motion correction; 6-mm FWHM smoothing; signal normalization; transformation to Talairach space | AlphaSim; 1,000 Monte Carlo simulations | Voxelwise $p < .005$; $k >= 6$ voxels | AFNI | 1.5T GE MRI scanner |
| Blackwood et al. (2004) | schizophrenia | Discard first 4 volumes; motion correction; 6-mm FWHM smoothing; signal normalization; transformation to Talairach space | AlphaSim; 1,000 Monte Carlo simulations | Voxelwise $p < .005$ uncorrected; mapwise corrected $p < .05$ | AFNI | 1.5T GE MRI scanner |
| Blair et al. (2008) | social anxiety disorder | Discard first 4 volumes; realignment; unwarping; normalization to MNI EPI template (2-mm isotropic); 8-mm FWHM smoothing | AlphaSim; 10,000 simulations | Voxelwise $p < .005$ uncorrected; $k >= 141$ voxels; cluster FWE $p < .05$ | SPM8 | 1.5T GE Signa |

| | | | | | | |
|---|---|---|---|---|---|---|
| Blair et al. (2011) | social anxiety disorder | 100-s high-pass filter; 5-mm FWHM smoothing; skull stripping; registration; motion-outlier regression; ICA-based denoising when indicated | FSL cluster correction | $Z > 2.3$; whole-brain cluster probability $p < .05$ | FSL 5.0.9 | 3T Siemens Tim Trio |
| Hashimoto et al. (2017) | autism spectrum disorder | Discard first 3 volumes/run; slice-timing correction; realignment; T1 coregistration; MNI normalization; 8-mm FWHM smoothing | FDR outside a priori MFG; uncorrected within MFG | MFG: $p < .001$ uncorrected, $k >= 5$ voxels; elsewhere FDR $p < .05$ | SPM5 | 1.5T GE Signa |
| Jimenez et al. (2018) | schizophrenia | Timing and motion correction; MNI registration; resampling to 3-mm isotropic voxels; Gaussian spatial filtering; white-matter/noise signal removal | AlphaSim whole-brain correction | Group comparisons $p < .025$; $k > 71$ voxels; corrected $p < .05$ | SPM algorithms; BioImage Suite; in-house R | 3T Siemens scanner |
| Lemogne et al. (2009) | major depressive disorder | Slice-timing correction; realignment; coregistration; segmentation; MNI normalization; 8-mm FWHM smoothing; high-pass temporal filtering | FDR; FDR SVC for a priori regions | FDR $p < .05$; $k >= 25$ voxels; SVC extent NR | SPM5 | 3T GE Signa |
| Liu et al. (2014) | schizophrenia | Realignment to first volume; motion inspection; MNI normalization (2-mm isotropic); 8-mm FWHM smoothing; high-pass filtering | Monte Carlo cluster correction; 10,000 iterations | Voxelwise $p < .001$; $k >= 80$ voxels; corrected $p = .01$ | SPM8 | 3T Siemens Magnetom |

| | | | | | | |
|---|---|---|---|---|---|---|
| Lombardo et al. (2010) | autism spectrum disorder | Realignment to first image; MNI normalization; 6-mm FWHM smoothing; 128-s high-pass filter | Cluster-level FWE | Voxelwise $p < .005$ uncorrected; cluster FWE $p < .05$ | SPM8 | 3T Philips Achieva |
| Makowski et al. (2016) | schizophrenia | Susceptibility-by-motion correction; realignment; MNI normalization; 6-mm FWHM smoothing; 128-s high-pass filter | Uncorrected extent threshold; SVC for hypothesized ROIs | Voxelwise $p < .005$ uncorrected; $k >= 50$ voxels; ROI SVC | SPM2 | 4T Bruker MedSpec |
| McAdams et al. (2016) | anorexia nervosa | Slice-timing correction; field-map unwarping; coregistration; unified segmentation/MNI normalization; 8-mm FWHM smoothing | Voxel-level FWE within task-effect voxels | Task effect: whole-brain voxel FWE $p < .05$; group interaction: FWE $p < .05$ within task-effect mask | SPM8 | 3T Siemens Trio |
| Murphy et al. (2010) | schizophrenia | Realignment; stereotaxic normalization; 8-mm FWHM smoothing; 1/128-Hz high-pass filter | AlphaSim; 1,000 simulations | Voxelwise $p = .001$; $k >= 16$ voxels; corrected $p < .05$ | SPM5 | 3T Philips scanner |
| Pankow et al. (2016) | schizophrenia | Motion correction; slice-timing correction; 6-mm FWHM smoothing; linear-trend removal and 0.02-Hz high-pass filtering; coregistration; Talairach transformation | Combined voxel- and extent-threshold rule | Voxelwise $p < .005$ uncorrected; $k >= 8$ native voxels | BrainVoyager | 3T Siemens Trio |

| | | | | | | |
|---|---|---|---|---|---|---|
| Pauly et al. (2014) | schizophrenia | Detailed preprocessing referred to an online supplement; main text reports 128-s high-pass filtering | Cluster-level FWE | Cluster-defining $t$ = 2.4; two-tailed cluster FWE $p < .05$ | SPM5 | 3T Philips Achieva |
| Sarsam et al. (2013) | major depressive disorder | Slice-timing correction; realignment; coregistration; MNI normalization; 8-mm FWHM smoothing | AlphaSim whole-brain correction | Voxelwise $p < .01$; whole-brain corrected $p < .01$, two-tailed | SPM5 | 2T GE/Elscint Prestige |
| Shad et al. (2012) | schizophrenia | Manual reorientation; slice-timing correction; realignment; coregistration; MNI normalization; 10-mm FWHM smoothing; high-pass filtering based on 1.1× longest intertrial interval | FWE for overall task; uncorrected ROI group tests | Group tests $p < .001$ uncorrected; $k >= 10$ voxels | SPM8 | 3T Philips Intera |
| Tan et al. (2015) | schizophrenia | Slice-timing correction; realignment; anatomical coregistration; MNI normalization; 10-mm FWHM smoothing; high-pass filtering based on 1.1× longest intertrial interval | Voxel-level FWE; cluster-level FWE within ROIs | Activation: voxel FWE $p < .05$; group effects: $k >= 10$ voxels and ROI cluster FWE $p < .05$ | SPM8 | 3T Philips Intera |
| van der Meer et al. (2013) | schizophrenia | MNI normalization; motion correction; 4-mm FWHM spatial smoothing | Uncorrected whole-brain and post-hoc/PPI analyses | Voxelwise $p < .001$ uncorrected; $k > 10$ voxels | SPM5 | 1.5T Siemens Symphony |

| | | | | | | |
|---|---|---|---|---|---|---|
| Zhang et al. (2015) | bipolar disorder and schizophrenia | Slice-timing correction; realignment; anatomical coregistration; DARTEL-based MNI normalization; 8-mm FWHM smoothing | Cluster-level FWE | Voxelwise $p < .001$; cluster FWE $p < .05$ | SPM12 | 3T Philips Ingenia |
| Yoshimura et al. (2010) | major depressive disorder | Realignment to first volume; MNI normalization; 6-mm FWHM smoothing | Uncorrected group comparison; Bonferroni-corrected ROI follow-up | Group comparisons: $p < .001$ uncorrected, $k >= 20$ voxels; ROI $p = .0125$ | SPM5 | 3T Philips scanner |
| Porta-Casteràs et al. (2023) | bipolar disorder | Slice-timing correction; rigid-body realignment; anatomical coregistration; unified segmentation/normalization to MNI; 8-mm FWHM smoothing | Whole-brain cluster-level FDR; additional ROI analyses | Voxelwise $p < .01$; whole-brain cluster FDR $p < .05$ | SPM8 | 3T Siemens Magnetom Skyra |

| | | | | | | |
|---|---|---|---|---|---|---|
| McAdams and Krawczyk (2014) | anorexia nervosa | Realignment; slice-timing correction; T1 bias correction and segmentation; nonlinear MNI normalization to 3-mm voxels; 8-mm FWHM smoothing; motion and framewise-displacement regressors | Whole-brain TFCE with FWE correction | Whole-brain TFCE FWE $p < .05$; no cluster-forming threshold | SPM12 | 3T GE Signa |
| Komulainen et al. (2018) | major depressive disorder | Slice-timing correction; realignment; MNI normalization to 3-mm voxels; Gaussian spatial smoothing; six motion parameters | Frontal-mask SVC with cluster-level FWE | Initial voxelwise $p < .005$; frontal-mask cluster FWE $p < .05$ | SPM8 | 3T Siemens Magnetom Trio TIM |
| Dixon et al. (2022) | social anxiety disorder | Slice-acquisition-time correction; realignment; anatomical coregistration; nonlinear MNI normalization; 6-mm FWHM smoothing | Uncorrected extent threshold | Voxelwise $p < .001$ uncorrected; $k >= 20$ voxels | SPM12 | 3T Siemens Tim Trio |
| Li et al. (2017) | major depressive disorder | Despiking; slice-timing correction; motion correction; brain extraction; functional–structural coregistration; linear/nonlinear MNI warping; 5-mm FWHM smoothing; grand-median intensity rescaling | AFNI 3dClustSim; separate SVC mask | Voxelwise $p < .001$; $k >= 30$ voxels; corrected cluster $p < .05$ | AFNI; FSL 5.0.2 | 3T Siemens Tim Trio |
| Hur et al. (2021) | social anxiety disorder | NR in sufficient detail in the supplied article | ROI/SVC; exploratory whole-brain FDR | Voxelwise $p < .005$ uncorrected; SVC FWE $p < .05$; whole-brain FDR $p < .05$ was null | NR | 4T Varian Unity Inova |
| Beeney et al. (2016) | personality disorder | Motion correction; temporal-drift correction; intensity normalization; 6-mm FWHM spatial smoothing; removal of global | Monte Carlo cluster correction | Within groups $p < .005$; between groups $p < .05$; | AFNI; FreeSurfer; | 3T Siemens TIM Trio |

| | | | | | | |
|---|---|---|---|---|---|---|
| | | intensity variation; surface-based normalization | in predefined cortical territories | cluster-corrected $p < .05$ | SPM2 (between-group analysis) | |
| Bluhm et al. (2012) | post-traumatic stress disorder | Field-map correction; discard first 10 TRs; motion correction and censoring; slice-timing correction; 6-mm FWHM smoothing; linear-trend removal | AlphaSim extent correction | Voxelwise $p < .001$; minimum cluster volume 384 $mm^3$ | AFNI 2.56 | 3T GE Signa EXCITE |
| Holt et al. (2011) | schizophrenia | Realignment; MNI normalization; 8-mm FWHM smoothing; 128-s high-pass filter; motion parameters entered as regressors | Uncorrected extent threshold | Voxelwise $p < .001$ uncorrected; $k >= 10$ voxels | SPM8 | 3T GE Signa Excite |
| Kennedy and Courchesne (2008) | autism spectrum disorder | Realignment; unwarping; T1 coregistration; nonlinear normalization using segmentation parameters (2-mm isotropic); 6-mm FWHM smoothing | Voxel-level FWE within a priori SCC ROI | Display $p = .005$ uncorrected; ROI voxel FWE $p = .05$; whole-brain FWE analysis null | SPM8 | 3T Philips Achieva |
| Komeda et al. (2015) | autism spectrum disorder | Realignment; coregistration; MNI normalization; 8-mm FWHM smoothing; 128-s high-pass filter | Cluster-level FWE | Cluster-defining $t = 2.4$; two-tailed cluster FWE $p < .05$ | SPM5 | 3T Philips Achieva |

| | | | | | | |
|---|---|---|---|---|---|---|
| Lythe et al. (2022) | bipolar disorder | Discard first 7 images; despiking; motion correction; Talairach normalization; 6-mm FWHM smoothing | Monte Carlo cluster correction within network templates | Voxelwise $p < .01$; network-specific extent; corrected $p < .05$ | AFNI | 3T GE Signa Excite |

### S8. Supplementary results: Regional distribution and direction of significant direct group comparison in self-reflection contrast effects in the systematic review and meta-analyses.

**Table S4.** Regional distribution and direction of significant direct group comparison in self-referential contrast effects.

| Brain region | Higher activation contrast ↑ | Lower activation contrast ↓ |
|---|---|---|
| **A. Self > other** | | |
| Medial frontal / prefrontal cortex | Blair et al. (2008); Sarsam et al. (2013) | Bedford et al. (2012); Pankow et al. (2015) |
| Anterior cingulate cortex | - | Blackwood et al. (2004) |
| Posterior cingulate cortex | Blackwood et al. (2004) | Bedford et al. (2012); Makowski et al. (2016) |
| Middle cingulate cortex | - | Makowski et al. (2016) |
| Precuneus | - | McAdams et al. (2014) |
| Inferior frontal / ventrolateral cortex | Liu et al. (2014) | - |
| Lateral parietal cortex / TPJ | Fuentes-Claramonte et al. (2020) | Hashimoto et al. (2017); Makowski et al. (2016) |
| Right superior/posterior temporal cortex | Liu et al. (2014) | - |
| Left inferior temporal gyrus | - | Liu et al. (2014) |
| Parahippocampal / amygdala region | Blair et al. (2008) | - |
| Precentral / postcentral cortex | Shad et al. (2012) | - |
| Lingual / visual cortex | Shad et al. (2012) | - |

| B. Self > control | | |
|---|---|---|
| Medial frontal / prefrontal cortex | Lemogne et al. (2009); Porta-Casteràs et al. (2023) | Pankow et al. (2015); Tan et al. (2015); Bluhm et al. (2012) |
| Anterior cingulate cortex | - | Tan et al. (2015) |
| Medial orbitofrontal cortex | - | Holt et al. (2011) |
| Posterior cingulate cortex | Holt et al. (2011) | van der Meer et al. (2013) |
| Middle cingulate cortex | Holt et al. (2011) | - |
| Precuneus | - | Murphy et al. (2010) |
| Middle / superior lateral frontal cortex | Fuentes-Claramonte et al. (2020); Furuichi et al. (2019) | Pankow et al. (2015) |
| Inferior frontal / ventrolateral cortex | Lemogne et al. (2009) | - |
| Lateral parietal cortex / TPJ | Furuichi et al. (2019) | Murphy et al. (2010) |
| Temporal cortex | - | Pauly et al. (2014) |
| Lingual / visual cortex | - | Murphy et al. (2010) |

### S9. Supplementary results: Experiments with reported foci within significant ALE clusters.

**Table S6.** Experiments with reported foci within significant ALE clusters

| Significant cluster | Contributing experiments | Foci within cluster | Contributing experiments (n/N) |
|---|---|---|---|
| **Direction-agnostic single ALE: Both directions pooled; [self>other]** | | | |
| R anterior cingulate cortex (x,y,z: 8, 38, 0) | Bedford et al. (2012) - schizophrenia<br>Blackwood et al. (2004) - schizophrenia<br>Pankow et al. (2015) - schizophrenia | 3 | 3/13(23.1%) |
| **Direction-agnostic single ALE: Both directions pooled; [self>control]** | | | |
| L middle frontal gyrus (x,y,z: −44, 16, 40) | Fuentes-Claramonte et al. (2020) -schizophrenia<br>Pankow et al. (2015) - schizophrenia<br>Lemogne et al. (2009) - depression | 3 | 3/12(25.0%) |
| **Single ALE: $\text{Patients}_{[self>other]}<\text{controls}_{[self>other]}$** | | | |
| L anterior cingulate cortex (x,y,z: −6, 46, −4) | Lombardo et al. (2010) - autism spectrum disorder<br>Pankow et al. (2015) - schizophrenia | 2 | 2/8 (25.0%) |
| **Single ALE: $\text{Patients}_{[self>control]}<\text{controls}_{[self>control]}$** | | | |
| L middle frontal gyrus (x,y,z: −44, 16, 40) | Fuentes-Claramonte et al. (2020) -schizophrenia<br>Pankow et al. (2015) - schizophrenia | 3 | 2/9 (22.2%) |

***Note.*** FocusCounter was used to identify experiments with one or more reported peak coordinates located within each significant ALE cluster. Foci within cluster denotes the total number of reported peak coordinates located within the cluster and may therefore exceed the number of contributing experiments. In *n/N*, *n* is the number of experiments containing at least one focus within the cluster, and *N* is the total number of experiments included in the corresponding ALE analysis. Peak coordinates are reported in MNI space.

**S10. Supplementary methods: Exploratory ALE subtraction analyses.**

We performed exploratory ALE subtraction analyses to determine whether the spatial convergence of patient–control differences varied between the self-versus-control and self-versus-other task-contrast categories. This question differed from that addressed by the individual ALE analyses: rather than testing whether coordinates converged within a given dataset, the subtraction analyses tested whether the spatial distributions of convergence differed between two sets of experiments (Laird et al., 2005).

The analyses were conducted separately for the two directions of the patient–control difference. First, the ALE map derived from experiments reporting lower self-versus-control contrast effects in psychiatric patients than in healthy controls was compared with the ALE map derived from experiments reporting lower self-versus-other contrast effects in psychiatric patients than in healthy controls. These datasets contained 9 and 8 experiments, respectively. Second, the corresponding comparison was conducted for higher patient–control contrast effects, for which the self-versus-control and self-versus-other datasets contained 3 and 8 experiments, respectively.

The subtraction analyses were implemented using NiMARE's ALESubtraction procedure. Experiments were randomly reassigned between the two datasets 10,000 times to generate an empirical null distribution of between-map differences. The resulting subtraction maps were evaluated using two-sided tests, thereby examining both greater convergence for self-versus-control than for self-versus-other and greater convergence for self-versus-other than for self-versus-control. Maps were thresholded at a voxel-level threshold of $p < .001$ (uncorrected), together with a minimum cluster extent of 25 contiguous $2 \times 2 \times 2$ mm voxels (200 $mm^3$).

Neither analysis identified a suprathreshold cluster. For lower patient–control contrast effects, no significant difference in spatial convergence was observed between the self-versus-control and self-versus-other datasets in either subtraction direction. Similarly, for higher patient–control contrast effects, no significant difference was detected between the two task-contrast categories in either subtraction direction.

### S11. Supplementary methods: Exploratory functional decoding.

To understand psychological process associated with the brain region identified by the ALE analyses, we conducted meta-analytic functional decoding (Smith et al., 2009). Specifically, we employed unthresholded $z$-value maps from the ALE results to retain the full spatial distribution of signal intensities, as recommended for continuous decoding approaches (https://nimare.readthedocs.io/en/stable/decoding.html). Functional decoding was then performed using the CorrelationDecoder module of the NiMARE Python package (version 0.20.0; Salo et al., 2023). This approach estimates psychological relevance by computing voxel-wise Pearson correlations between the input ALE map and term-based meta-analytic activation maps derived from Neurosynth (Yarkoni et al., 2011), a large-scale database that aggregates activation coordinates and term-based annotations from 14,371 published fMRI studies.

To restrict the list of terms to those that involve psychological processes, we filtered the Neurosynth terms using the Cognitive Atlas ontology (Poldrack et al., 2011), yielding 133 terms spanning broad and specific domains of cognitive processes, behavior types, and emotional states.

For each decoded map, the 10 terms with the highest positive correlations coefficients were retained for visualization. The results were interpreted descriptively as spatial similarities between the ALE and term-associated maps, rather than as evidence that psychological processes were selectively impaired or that the identified regions were specific to those processes.
